# Systemic and targeted activation of Nrf2 reverses doxorubicin-induced cognitive impairments and sensorimotor deficits in mice

**DOI:** 10.1101/2024.06.10.598291

**Authors:** Anand Kumar Singh, David Ruiz, Mohd Sami Ur Rasheed, Thomas D. Avery, Dion J. L. Turner, Andrew D. Abell, Peter M. Grace

## Abstract

While cancer survivorship has increased due to advances in treatments, chemotherapy often carries long-lived neurotoxic side effects which reduce quality of life. Commonly affected domains include memory, executive function, attention, processing speed and sensorimotor function, colloquially known as chemotherapy-induced cognitive impairment (CICI) or “chemobrain”. Oxidative stress and neuroimmune signaling in the brain have been mechanistically linked to the deleterious effects of chemotherapy on cognition and sensorimotor function. With this in mind, we tested if activation of the master regulator of antioxidant response nuclear factor E2-related factor 2 (Nrf2) alleviates cognitive and sensorimotor impairments induced by doxorubicin. The FDA-approved systemic Nrf2 activator, diroximel fumarate (DRF) was used, along with our recently developed prodrug **1c** which has the advantage of specifically releasing monomethyl fumarate at sites of oxidative stress. DRF and **1c** both reversed doxorubicin-induced deficits in executive function, spatial and working memory, as well as decrements in fine motor coordination and grip strength, across both male and female mice. Both treatments reversed doxorubicin-induced loss of synaptic proteins and microglia phenotypic transition in the hippocampus. Doxorubicin-induced myelin damage in the corpus callosum was reversed by both Nrf2 activators. These results demonstrate the therapeutic potential of Nrf2 activators to reverse doxorubicin-induced cognitive impairments, motor incoordination, and associated structural and phenotypic changes in the brain. The localized release of monomethyl fumarate by **1c** has the potential to diminish unwanted effects of fumarates while retaining efficacy.

## Introduction

Cancer survivorship has increased due to the successes of both early detection and effective treatments. As of 2022, there were more than 18 million cancer survivors in the US alone, and it is projected that these numbers will increase to 22.5 million by 2032 (Miller, Nogueira et al. 2022). Unfortunately, treatment of cancers with chemotherapy is often accompanied by long-lived neurotoxic side effects (Bolton and Isaacs 2018). Commonly affected domains include memory, executive function, attention, processing speed, and sensorimotor function, known as chemotherapy-induced cognitive impairment (CICI), or colloquially ‘chemobrain’. There are no FDA-approved treatments for these deficits which persist years into survivorship, dramatically reducing quality of life (Bolton and Isaacs 2018, Binarelli, Duivon et al. 2023, Ossorio-Salazar and D’Hooge 2023).

Several groups have identified chemotherapy-induced abnormalities in hippocampus and prefrontal cortex as underlying cognitive impairment. These include reduced numbers of neuronal spines and dendrites, lower neurogenesis, impaired myelination and persistent dysregulation of oligodendrocyte, astrocyte, and microglial function which are all crucial for health and plasticity of the central nervous system (Gibson, Nagaraja et al. 2019, Nguyen and Ehrlich 2020, Liu, Reiken et al. 2023). The mechanisms by which oncology drugs exert these cellular effects are varied due to differing therapeutic regimens and mechanisms of action between classes. However, common mechanisms underpinning cognitive dysfunction have been identified (Nguyen and Ehrlich 2020). Several oncology drugs, including doxorubicin, used to treat breast cancer, enhance production of reactive oxygen species (ROS), leading to oxidative stress. This in turn contributes to mitochondrial dysfunction diminished DNA repair capacity, and chronic inflammation (Maynard, Fang et al. 2015, Ren, St Clair et al. 2017, Keeney, Ren et al. 2018).

A recently developed chemobrain mouse model using doxorubicin shows long-term cognitive impairments associated with synapse damage in the hippocampus (McAlpin, Mahalingam et al. 2022). These neurotoxic effects occur indiscriminately, often affecting other brain regions and impairing function (Chiang, Seua et al. 2020). For example, cancer survivors also exhibit sensorimotor deficits, showing decreased fine motor dexterity over time which often persists for months (Hile, Fitzgerald et al. 2010, Grusdat, Stauber et al. 2022). The mechanism of the decline in the fine motor skill is largely unknown despite CNS imaging data of breast cancer patients treated with high doses of carmustine, cyclophosphamide, and cisplatin suggesting decrease in the white matter (Brown, Stemmer et al. 1998). Higher loads of ROS result in a decrease in white matter by altering the dynamics of myelin producing cells. Differentiating oligodendrocyte precursor cells are most vulnerable against increased oxidative stress (Dracheva, Davis et al. 2006, Maas, Valles et al. 2017).

Despite widespread recognition that oxidative stress is a key mechanistic node, ROS have yet to be therapeutically targeted to treat CICI or sensorimotor deficits. However, activation of nuclear factor E2-related factor 2 (Nrf2) has emerged as a new strategy to restore redox homeostasis (Cuadrado, Rojo et al. 2019, Dinkova-Kostova and Copple 2023). Nrf2 is an endogenous master regulator of genes involved in antioxidant metabolism and anti-inflammation (Hayes and Dinkova-Kostova 2014, Cuadrado, Rojo et al. 2019). Under basal conditions, cytoplasmic Nrf2 is bound with Kelch-like ECH-associated protein 1 (Keap1), which targets the complex for proteasomal degradation (Kobayashi, Kang et al. 2004). Electrophilic pharmacological agents such as monomethyl fumarate and sulforaphane bind to Keap1, disrupting the complex and stabilizing Nrf2 to allow translocation to the nucleus (Suzuki and Yamamoto 2015, Yamamoto, Kensler et al. 2018). In the nucleus, Nrf2 binds to antioxidant response elements and induces the transcription of antioxidant genes (Cuadrado, Manda et al. 2018). The Nrf2 pathway is therefore a potential therapeutic target for CICI, with its activation predicted to resolve underlying oxidative stress and neuroinflammation.

Fumarates drugs are effective Nrf2 activators in this context with clinical efficacy in treating recurring remitting multiple sclerosis, and preclinical efficacy in numerous other neurological diseases and disorders (Rostami-Yazdi, Clement et al. 2010, Naismith, Wolinsky et al. 2020, Hoogendoorn, Avery et al. 2021). Diroximel fumarate (DRF) is a prodrug which systemically releases the therapeutic metabolite and Nrf2 activator, monomethyl fumarate (MMF) (Hoogendoorn, Avery et al. 2021). Since Nrf2 is ubiquitously expressed in all the cells across the body, off-target effects by systemic Nrf2 activators result (Cada, Levien et al. 2013). We recently developed a targeted prodrug (compound **1c**) which releases MMF on exposure to ROS elevated at sites of pathology (Avery, Li et al. 2021). This then reduces the dose-limiting adverse effects of fumarates and of global Nrf2 activation (Cuadrado, Rojo et al. 2019, Avery, Li et al. 2021).

In the current study, we tested the capacity of systemic (DRF) and locally delivered (compound **1c**) MMF to reverse cognitive impairments induced by doxorubicin, and the associated decrease in post-synaptic protein and changes in microglia morphology. We also examined the influence of DRF and **1c** on doxorubicin-induced sensorimotor deficits and associated loss of myelin in the corpus callosum of the mouse brain.

## Methods

### Animals

Male and female C57BL/6J mice (8-12 weeks) were purchased from Jackson Laboratory and housed in MD Anderson Cancer Center animal facility on a 12/12 h reverse light cycle at 22 ± 2°C with water and food available *ad libitum*. All experimental procedures were consistent with the National Institute of Health Guidelines for the Care and Use of Laboratory Animals and were approved by the MD Anderson Cancer Center Animal Care and Use Committee. Mice were randomly assigned to experimental groups and all behavioral tests were performed by an investigator blinded to group assignments. Before the behavioral testing, each mouse was handled by the experimenter, 1-2 min for 3 days in the test area. Video recording of cognitive behavior and beam walk test was done under red light using a Bell & Howell, Infrared Night Vision Camcorder.

### Drugs

Doxorubicin (5 mg/kg/week, Pfizer, New York, NY) was diluted in sterile phosphate-buffered saline (PBS) and treated for 4 weeks, one injection intraperitoneally each week. Diroximel fumarate (DRF, 89 mg/kg, oral; MedChemExpress) or **1c** (60 mg/kg, oral; synthesized by T. Avery) was suspended in PBS containing 2% methylcellulose and administered daily, starting one week after doxorubicin treatment and continuing throughout the cognitive and sensorimotor behavioral tests. These doses were selected based on our prior studies (Avery, Li et al. 2021). PBS was used as the vehicle control for cisplatin, 2% methylcellulose solution for DRF and **1c**. Two weeks after the completion of doxorubicin treatment (and after one week of DRF or **1c** treatment), mice were tested for cognitive behavior and sensorimotor function followed by tissue collection for biochemical assays. The experimental timeline is presented in **Fig. S1**.

### Puzzle box test (PBT)

The puzzle box test for executive functioning was performed as described previously (Chiu, Maj et al. 2017, McAlpin, Mahalingam et al. 2022, Singh, Mahalingam et al. 2022). Mice were placed individually into a brightly illuminated arena from which they could escape to a dark goal box by a tunnel. Time to reach the goal box was assessed when the tunnel was open (easy trials 1-4, day 1-2), filled with bedding (intermediate trials 5-7, day 2-3), or blocked with a plug (difficult trials 8-11, day 3-4).

### Novel object place recognition test (NOPRT)

The NOPRT for short-term memory and place recognition was performed as described (Chiu, Maj et al. 2017, McAlpin, Mahalingam et al. 2022, Singh, Mahalingam et al. 2022). During training, mice were introduced to two identical objects placed on one side of a rectangular arena for 5 min. After 1 h, mice were returned to the arena containing one familiar object at the same location as in the training session, and one novel object (Rubik’s cube) placed in a novel location. Interaction times with each of the objects were recorded for 5 min and analyzed with EthoVision XT 10.1 video tracking software (Noldus Information Technology Inc., Leesburg, VA). The discrimination index was calculated as (TNovel – TFamiliar)/ (TNovel + TFamiliar).

### Sensorimotor function

In the beam walk test for sensorimotor function, mice were trained on 3 consecutive days to cross a wide rectangular beam (3 trials), followed by a narrow rectangular or round beam until control mice successfully crossed the beam. The crossing time was determined in 2 videotaped trials analyzed by investigators blinded to treatment (Singh, Mahalingam et al. 2022).

### Grip strength

Optimal communication between motor neurons and muscle fibers is extremely important for maintenance of the neuromuscular function (Grinnell 1995). A Grip Strength Meter was used to measure fore and hind limb grip strength. As a mouse grasped the bar, the peak pull force in grams was recorded on a digital force transducer. Two trials were averaged and normalized with mouse weight.

### Immunohistochemistry

At the end of the behavioral analysis, mice were perfused transcardially with ice-cold PBS with 5 U/mL sodium heparin (Hospira). Brains were post-fixed in 4% PFA for 48 h and cryo-protected in sucrose. Coronal sections (8 or 20 µm) were blocked in 2% bovine serum albumin, 10% normal goat serum and 0.1% saponin in PBS followed by incubated with rabbit anti-PSD95 (1:1000; Abcam; AB18258) or rabbit anti-Iba1 (1:1000; Wako; 019-19741) diluted in antibody buffer (2% normal goat serum, 2% bovine serum albumin and 0.1% saponin in PBS) at 4 °C overnight. As a negative control, the primary antibody was omitted. Sections were then washed three times with PBS, followed by incubation with Alexa-488 goat anti-rabbit (1:500; Invitrogen; A21206) or Alexa-647 goat anti-rabbit (1:500; Invitrogen; A21245) at room temperature for 2 h. Fluorescence was visualized using the Nikon A1R Confocal Microscope (Nikon Instruments Inc., Melville, NY, USA) using a 40X objective. The mean intensity of PSD95 positive puncta was quantified in the CA1 of the hippocampus in three regions of interest using the spot detection feature of the Nikon NIS-Elements Advanced Research Software (Nikon Instruments Inc., Melville, NY, USA) (McAlpin, Mahalingam et al. 2022, Singh, Mahalingam et al. 2022). Sholl analysis of microglia was performed using the ImageJ simple neurite tracer module of the Neuroanatomy plugin and the area under the curve was quantified.

### Transmission electron microscopy (TEM)

For TEM analysis of myelin integrity, mice were anesthetized and transcardially perfused with PBS. One hemisphere of the brain was post-fixed in 2% glutaraldehyde plus 2% PFA in PBS at 4°C for at least a week. Small biopsy sample extracts about 1 mm in diameter and 2 mm in length were dissected out from the motor cortex. Fixed samples were processed at the High-Resolution Electron Microscopy Core at MD Anderson. Briefly, samples were washed in 0.1 M sodium cacodylate buffer and treated with cacodylate buffered tannic acid, post-fixed with 1% buffered osmium and stained in bloc with 0.1% Millipore-filtered uranyl acetate. Samples were then dehydrated in increasing concentrations of ethanol and infiltrated and embedded in LX-112 medium. Samples were polymerized in a 60 °C oven for approximately 3 days. Ultrathin sections were cut using a Leica Ultracut microtome and then stained with uranyl acetate and lead citrate in a Leica EM Stainer. Stained samples were examined in a JEM 1010 transmission electron microscope (JEOL USA, Inc, Peabody, MA) using an accelerating voltage of 80 kV. Digital images were obtained using an AMT imaging system (Advanced Microscopy Techniques Corp., Danvers, MA). Axons were analyzed for g-ratios calculated by dividing the internal diameter by outer diameter (diameter of axon/diameter of axon + myelin sheath) at 20,000x using ImageJ software. For each animal, around 200 axons were scored, and the mean g-ratio was analyzed.

### Statistical analysis

Data were analyzed using GraphPad Prism version 8.0.0 for Windows (GraphPad Software, San Diego, CA, USA). Error bars indicate SEM and statistical significance was assessed by t-test or by one-way or two-way ANOVA followed by two-tailed Tukey’s test for post hoc pairwise, multiple-comparisons. P<0.05 was considered statistically significant.

## Results

### Nrf2 activators reverse doxorubicin-induced cognitive impairments in mice

Here we have tested if DRF or compound **1c** would alleviate cognitive impairments induced by doxorubicin. DRF is an FDA-approved prodrug, which systemically distributes MMF, and consequently activates Nrf2 globally. Our groups have developed **1c**, a prodrug which releases MMF locally at sites of oxidative stress (Avery, Li et al. 2021). DRF or **1c** were administered daily to male and female mice, beginning one week post conclusion of doxorubicin treatment and continuing for 5 weeks (during the cognitive and sensorimotor behavioral tests) (**Fig. S1**). Doxorubicin significantly reduced performance during the difficult trials, with doxorubicin-treated male and female mice requiring more time than control mice to enter the dark compartment (**Fig. 1A-B**), consistent with previous results (McAlpin, Mahalingam et al. 2022). DRF and **1c** reversed these deficits, with performance similar to that of control mice (see **Fig. 1A-B**).

**Fig. 1:**
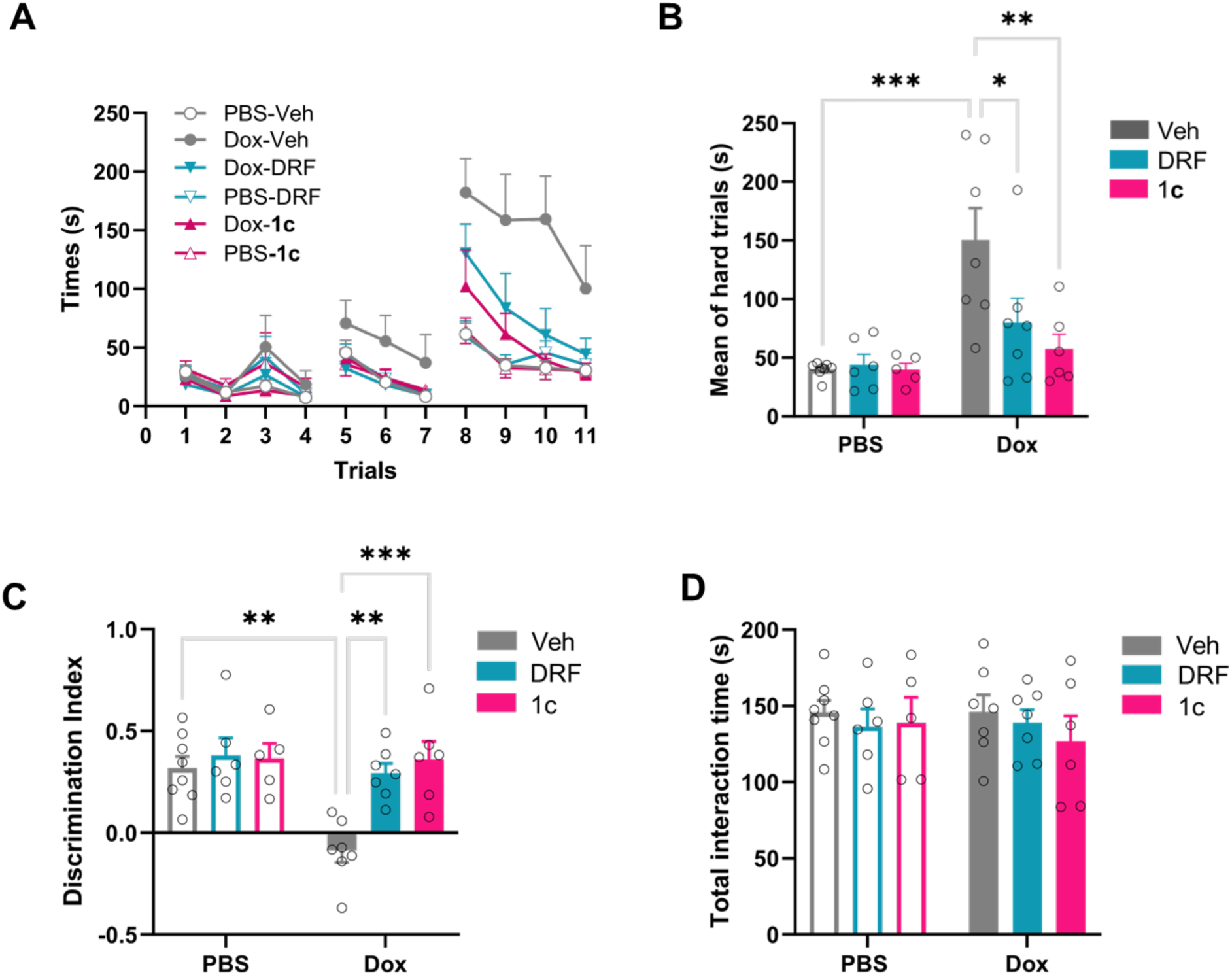
Global and local NRF2 activators reverses doxorubicin-induced cognitive impairments. (A-B) Influence of DRF and **1c** on doxorubicin-induced deficits in the executive function as assessed using the puzzle box test (PBT). (A) Time taken by mice to enter the dark chamber in all 11 trials (easy trials 1-4, intermediate trials 5-7 and hard trials 8-11). (B) Mean of hard trials in which mice have to remove a plug blocking the tunnel to the dark compartment to escape from the light compartment. Results are expressed as mean ± SEM; Two-way ANOVA with Tukey’s posthoc analysis. Chemotherapy x Nrf2 activator: F (2, 33) = 4.605, P = 0.0172; Chemotherapy: F (1, 33) = 16.14, P = 0.0003; Nrf2 activator: F (2, 33) = 4.330, P = 0.0214. (C) Effect of doxorubicin and Nrf2 activators on working memory in the novel object place recognition test (NOPRT). Preference for the novel object in the NOPRT is presented as the discrimination index (DI): (Tnovel-Tfamiliar)/ (Tnovel + Tfamiliar). Results are expressed as mean ± SEM; Two-way ANOVA with Tukey’s posthoc analysis. Chemotherapy x Nrf2 activator: F (2, 33) = 4.908, P = 0.0136; Chemotherapy: F (1, 33) = 8.569, P = 0.0062; Nrf2 activator: F (2, 33) = 8.304, P=0.0012. (D) Total interaction time in the NOPRT. Results are expressed as mean ± SEM; Two-way ANOVA with Tukey’s posthoc analysis. Chemotherapy x Nrf2 activator: F (2, 33) = 0.2034, P = 0.8170; Chemotherapy: F (1, 33) = 0.09525, P = 0.7595; Nrf2 activator: F (2, 33) = 0.6215, P=0.5433. *p < 0.05; **p < 0.01; ***p < 0.001. No significant sex effects were detected.

A novel object place recognition test (NOPRT) was performed to test working and spatial memory. Doxorubicin reduced the preference of male and female mice for the novel object, indicating memory deficits (**Fig. 1C**). In contrast, DRF and **1c** completely reversed the effect of doxorubicin on performance in the NOPRT (**Fig. 1C**). No differences were observed in the total interaction times between treatment groups, suggesting equal motivation for the exploration (**Fig. 1D**). These data collectively demonstrate that Nrf2 activators restore working and spatial memory and normalize executive function in doxorubicin-treated mice of both sexes.

### Global and local Nrf2 activation reverses doxorubicin-associated loss of post-synaptic protein in the CA1 region of the hippocampus

Chemotherapies are known to cause structural damage to synapse in CA1 of the hippocampus of mouse brain, which is associated with cognitive impairments (McAlpin, Mahalingam et al. 2022, Singh, Mahalingam et al. 2022). To evaluate changes in synaptic structure, we performed immunostaining of the post-synaptic protein PSD95 in the CA1 region of the hippocampus. Doxorubicin-treated mice had fewer PSD95 puncta compared to control (**Fig. 2A-B**). DRF or **1c** reversed these doxorubicin-induced decrements with intact PSD95 immunostaining and puncta number, suggesting either global or local Nrf2 activation reverses the doxorubicin-induced loss of post-synaptic protein (**Fig. 2A-B)**.

**Fig. 2:**
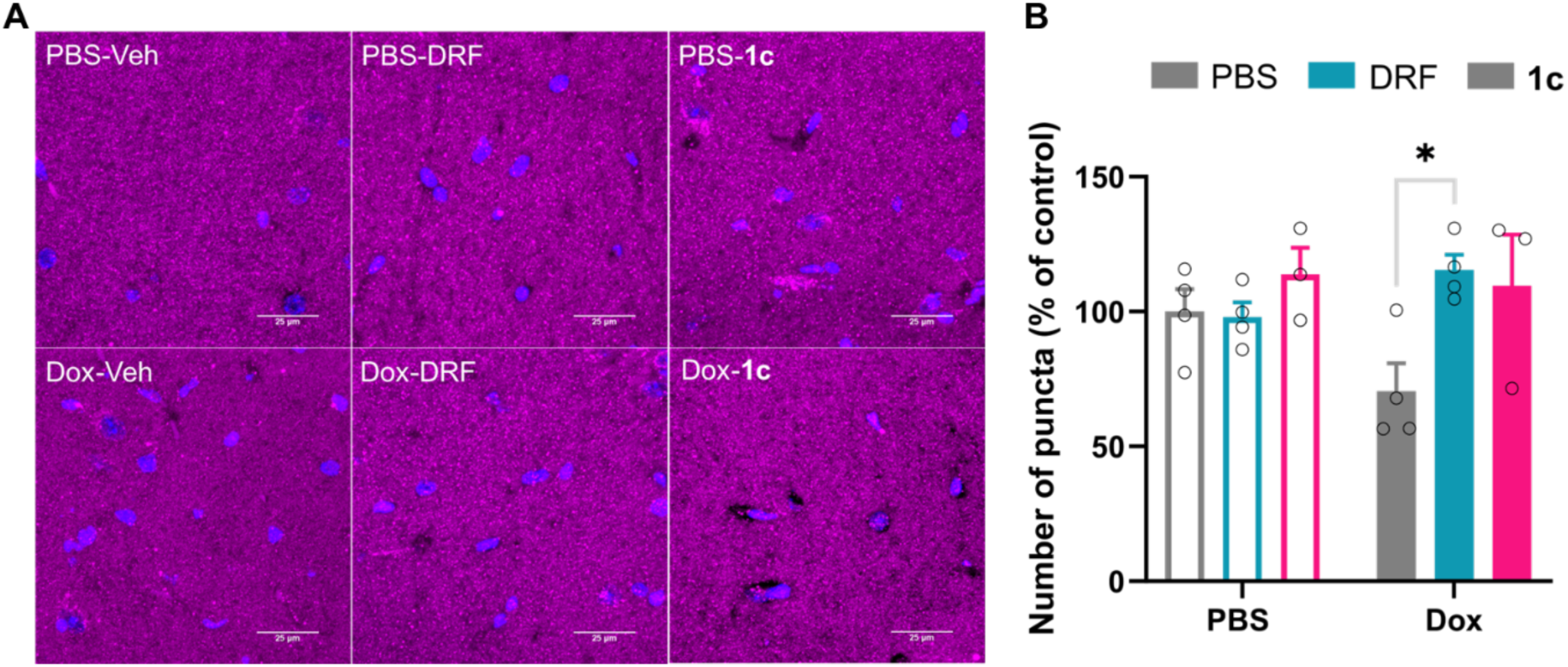
Doxorubicin-induced loss of post synaptic protein is reversed by Nrf2 activation. (A) Immunofluorescence images of the post-synaptic marker protein, PSD95 in the CA1 region visualized by confocal microscopy, scale bar 25 µm. (B) Number of puncta of the PSD95 protein are expressed as % of control. Results are expressed as mean ± SEM; Two-way ANOVA with Tukey’s post-hoc analysis. Chemotherapy x Nrf2 activator: F (2, 16) = 3.141, P = 0.0707; Chemotherapy: F (1, 16) = 0.4613, P = 0.5067; Nrf2 activator: F (2, 16) = 4.161, P = 0.0351. *p < 0.05.

### Nrf2 activators reverse doxorubicin-induced phenotypic change of microglia

Given the known antioxidant and anti-inflammatory effects of Nrf2 activators, we tested if DRF and **1c** reversed doxorubicin-induced morphological changes of microglia. The morphology of Iba1^+^ microglia was characterized using Sholl analysis (Hoogendoorn, Avery et al. 2021). Doxorubicin treatment decreased the branch level of microglia in the CA1 region of the hippocampus (**Fig. 3A**). Treatment of doxorubicin with either DRF or **1c** restored branching to the levels of controls (**Fig. 3A**). Sholl analysis revealed a robust decrease in the number of intersections in doxorubicin-treated mice compared to controls, indicating reduced microglia ramification (**Fig. 3B-C**). Treatment with DRF or **1c** reversed doxorubicin-induced decrease in the number of intersections, indicating that Nrf2 activators return morphology of microglia in CA1 to homeostasis (**Fig. 3D)**. Group differences in microglia morphology were determined using the area under the Sholl curve (AUSC). The AUSC was significantly smaller compared to control in the CA1 of doxorubicin-treated animals. The AUSC were higher for the mice receiving doxorubicin followed by DRF or 1c (**Fig. 3D**). There was no significant effect of DRF alone on the AUSC, but we observed higher AUSC in the **1c** alone treated group compared to control.

**Fig. 3:**
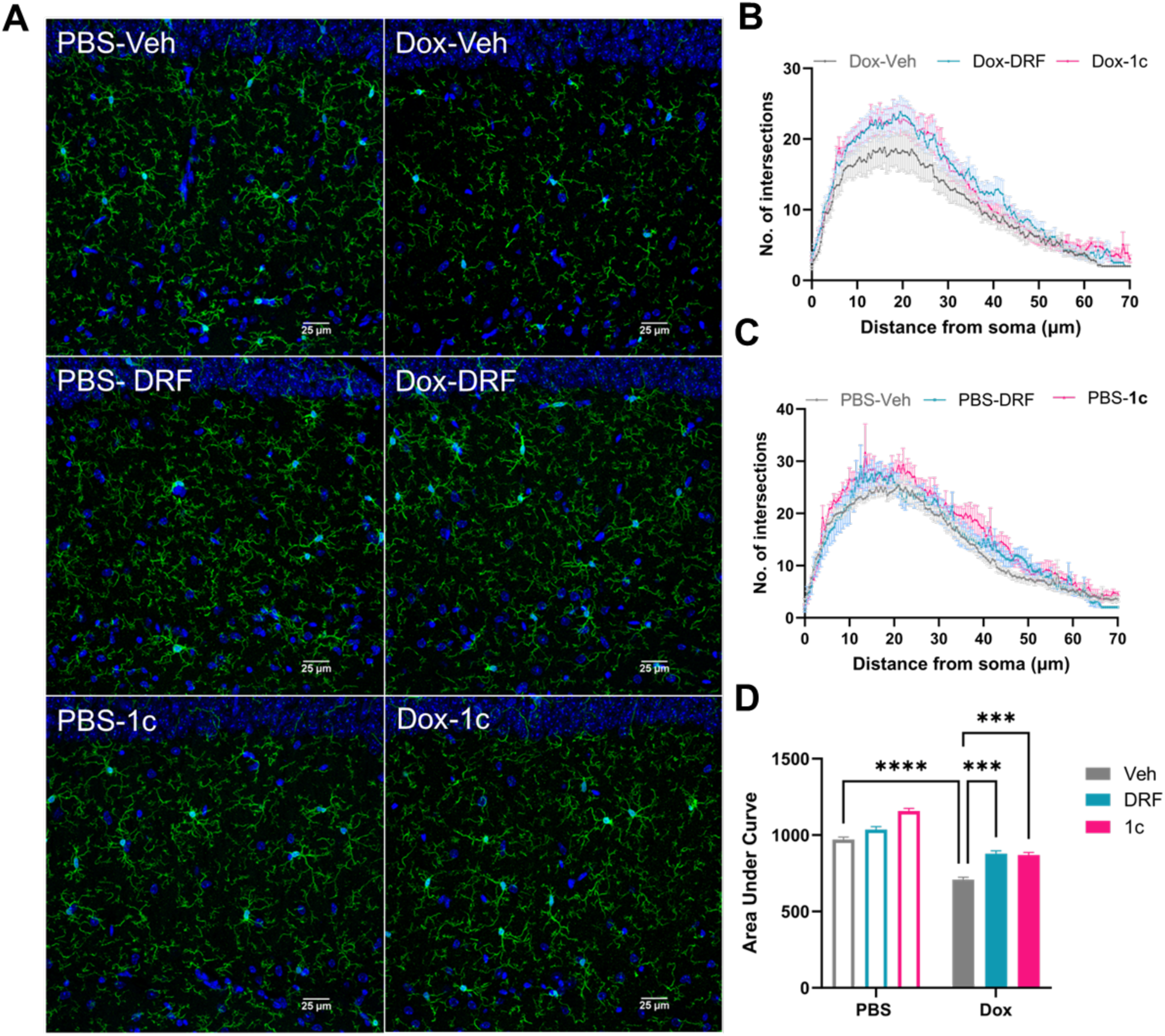
Nrf2 activators reverses doxorubicin-induced microglial phenotypic transition. (A) Confocal image showing Iba1 immunostaining of microglia. **(B-C)** Sholl analysis quantifies the number of projection intersections per concentric spherical shell beginning at the soma and distanced at a radius of 1 μm apart. **(D)** Quantification of the area under the Sholl curve from B. Results are expressed as mean ± SEM; Two-way ANOVA with Tukey’s posthoc analysis. Chemotherapy x Nrf2 activator: F (2, 11) = 6.749, P = 0.0122; Chemotherapy: F (1, 11) = 255.3, P<0.0001; Nrf2 activator: F (2, 11) = 52.70, P<0.0001. ***p < 0.001; ****p < 0.0001.

### Influence of Nrf2 activation on doxorubicin-mediated sensorimotor deficit and neuromuscular dysfunction

In addition to long-term cognitive impairments, chemotherapy treatment induces sensorimotor deficits (Hile, Fitzgerald et al. 2010, Chiang, Seua et al. 2020, Grusdat, Stauber et al. 2022). A beam walking test was carried out 4 weeks after the completion of doxorubicin treatment to measure sensorimotor function in mice. Mice treated with doxorubicin had an increased latency to cross the round beam (greatest difficulty), signifying deficits in fine sensorimotor function (**Fig. 4A**). The performances of mice treated with doxorubicin and DRF or **1c** were similar to controls (**Fig. 4A**). Neither doxorubicin, DRF, nor **1c** affected the time to cross wide and narrow flat beams, indicating that there was no difference in motivation to the subjected task (*Wide flat beam*: PBS-Veh 6.125 ±0.712 s; Dox-Veh 9.357±2.227 s; Dox-DRF 5.357±0.531 s; Dox-1c 6.750±1.039 s; PBS-DRF 6.000±0.548 s; PBS-1c 4.800±0.583 s and *Narrow flat beam*: PBS-Veh 11.437 ±1.814 s; Dox-Veh 11.857±1.383 s; Dox-DRF 7.714±0.778 s; Dox-1c 9.250±1.509s; PBS-DRF 10.000±1.658 s; PBS-1c 8.600±1.684 s). Thus, Nrf2 activators reversed doxorubicin-induced sensorimotor deficits in the of both sexes. (**Fig. 4A**).

**Fig. 4:**
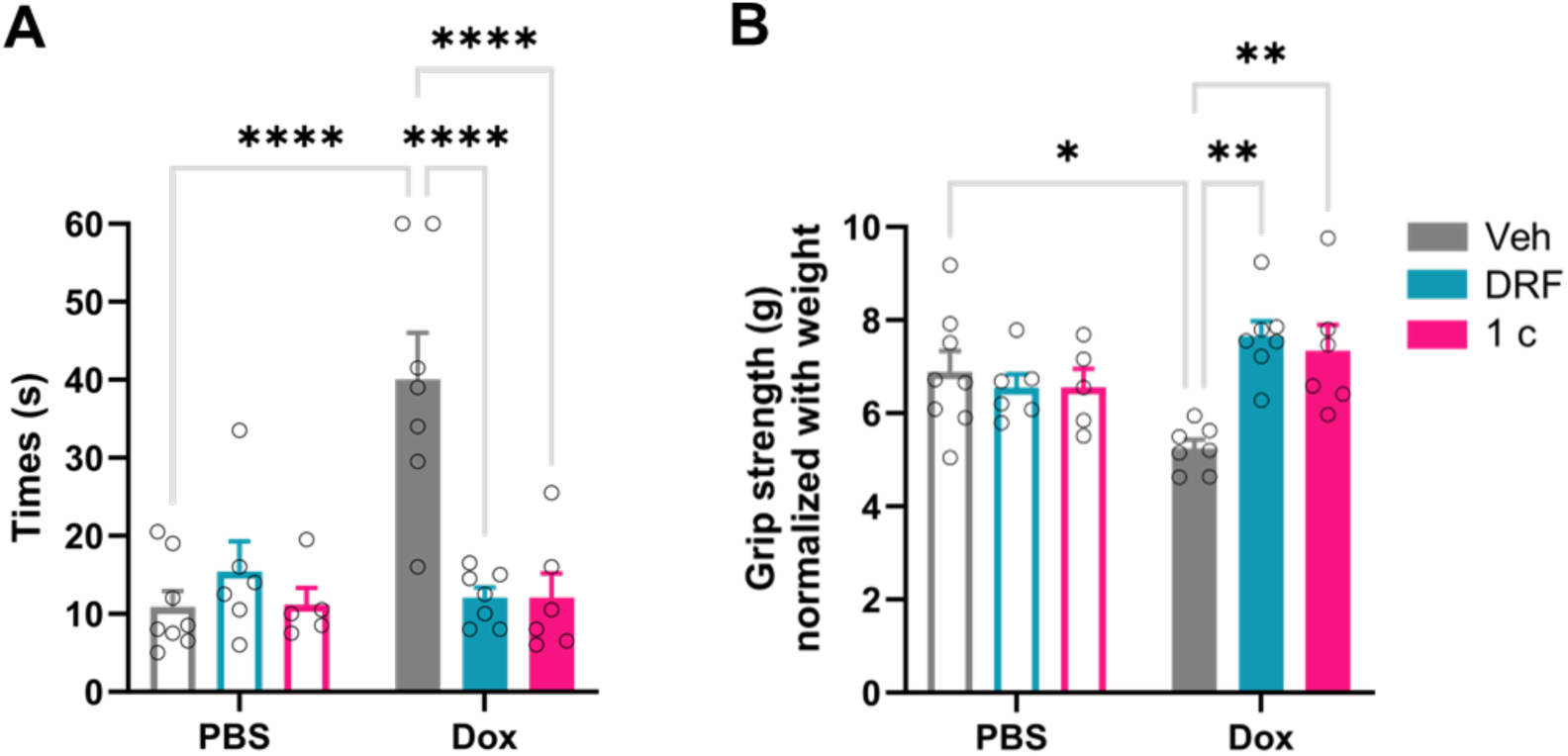
Influence of Nrf2 activation on doxorubicin-induced sensorimotor deficits and neuromuscular dysfunction. (A) The sensorimotor function measured by beam walking test. Three beams of different dimensions were used: wide flat, narrow flat and round. Data are expressed as the time it takes to walk the round beam (length 85 cm; diameter 6 mm). Results are expressed as mean ± SEM; Two-way ANOVA with Tukey’s posthoc analysis. Chemotherapy x Nrf2 activator: F (2, 33) = 13.43, P<0.0001; Chemotherapy: F (1, 33) = 9.338, P = 0.0044; Nrf2 activator: F (2, 33) = 9.279, P = 0.0006. (B) Grip strength of the mice presented normalized with weight. Results are expressed as mean ± SEM; Two-way ANOVA with Tukey’s posthoc analysis. Chemotherapy x Nrf2 activator: F (2, 33) = 7.742, P = 0.0018; Chemotherapy: F (1, 33) = 0.05845, P = 0.8105; Nrf2 activator: F (2, 33) = 4.341, P = 0.0212. *p < 0.05, **p < 0.01, ***p < 0.001, ****p < 0.0001.

Deficits in neuromuscular strength are also a common facet of sensorimotor impairment (Osaki, Morishita et al. 2022). Both fore– and hind-limb grip strength was reduced in doxorubicin-treated mice, compared to control mice (**Fig. 4B**). In contrast, mice treated with **1c** or DRF showed no doxorubicin-induced deficits in grip strength, demonstrating that Nrf2 activators rescued neuromuscular deficits in both sexes (**Fig. 4B**).

### Nrf2 activation reverses doxorubicin-induced myelin damage

Systemic chemotherapy is associated with progressive damage to myelin in animal models (Han, Yang et al. 2008, Chiang, Seua et al. 2020). Active myelination during adulthood is required for fine motor skills (McKenzie, Ohayon et al. 2014). Corpus callosum is the largest bundle of myelinated axons coming from cortical subregion from both side of hemisphere including the motor cortex (Wahl and Ziemann 2008). Transmission electron microscopy (TEM) was used to examine the impact of doxorubicin treatment on myelin ultrastructure in the corpus callosum of the motor cortex. The g-ratio was increased in doxorubicin-treated mice compared to control in each axonal diameter range, suggesting hypomyelination or loss of myelin layers due to treatment (**Fig. 5A-D**). G-ratios were normalized in mice treated with DRF or **1c**, indicating that Nrf2 activators reverse doxorubicin-induced myelin loss (**Fig. 5A-D**).

**Fig. 5:**
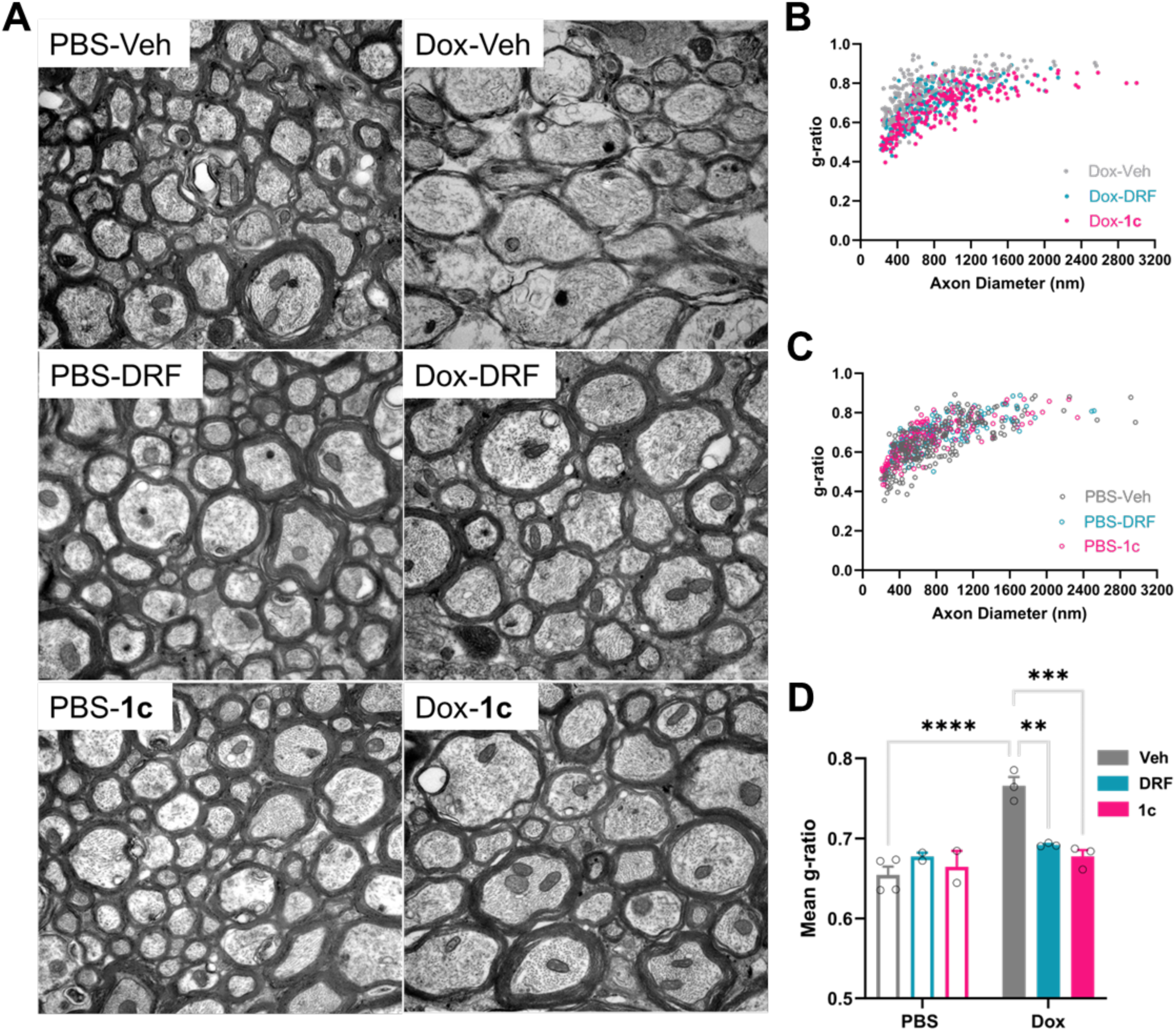
Nrf2 activators reversed doxorubicin-induced myelin damage. (A) Representative TEM images (20,000x) of corpus callosum axons with doxorubicin, DRF and **1c** treatment. (B-C) Scatterplots of g-ratio as a function of axon diameter measure quantified using ImageJ (D) Mean g-ratios for each mouse. Data are analyzed with two-way ANOVA (Tukey’s multiple comparisons test and results are expressed as mean ± SEM. Chemotherapy x Nrf2 activator: F (2, 11) = 15.67, P = 0.0006; Chemotherapy: F (1, 11) = 28.23, P=0.0002; Nrf2 activator: F (2, 11) = 7.545, P = 0.0087. **p < 0.01, ***p < 0.001, ****p < 0.0001.

## Discussion

Cancer patients undergoing chemotherapy with oncology drugs often suffer from long-term side effects, including a decline in cognitive function and fine motor dexterity which adversely affect the quality of life and functional independence (Brown, Stemmer et al. 1998, Wefel and Schagen 2012, Grusdat, Stauber et al. 2022). We show that Nrf2 activators reverse doxorubicin-induced cognitive and sensorimotor deficits in the mice of both sexes. This result shows that resolving oxidative stress induced by doxorubicin will alleviate deficits in cognitive and sensorimotor function. This normalization of behavior is associated with restored post-synaptic protein levels, resolution of microglia activation in CA1 of the hippocampus, and reversal of damage to the myelin sheath in the corpus callosum. These underlying structural and biochemical changes are reversed to a similar extent by either the systemic Nrf2 activator DRF, or local Nrf2 activator **1c** in mice of both sexes.

Oxidative stress in the brain has a significant impact on the cognitive function (Kandlur, Satyamoorthy et al. 2020). Neurons have high oxygen demands and enzymatic activity along with a large number of mitochondria, all of which are increased during long-term memory formation (Underwood, Redell et al. 2023). The higher metabolic demand under chemotherapy assault leads to enhanced release of oxidative species which damages synapse integrity and brain lipid especially myelin (Maas, Valles et al. 2017, Tonnies and Trushina 2017). After entering the body, doxorubicin is swiftly transformed into semiquinone and produces oxidative species after reacting with cytosolic, ER, and mitochondrial enzymes including NADH oxidase, NADPH cytochrome P450 reductase, xanthine oxidase and nitric oxide synthase (Deng, Kruger et al. 2007, Nithipongvanitch, Ittarat et al. 2007, Kong, Guo et al. 2022). A higher level of oxidative species has been reported in the plasma and brain the cancer patients and rodents treated with doxorubicin and were associated with cognitive decline (Faber, Coudray et al. 1995, Conklin 2004, Keeney, Ren et al. 2018).

A clinical study suggests a direct effect of doxorubicin in the brain (Du, Xia et al. 2020). Doxorubicin is reported to alter synaptic plasticity via reductions in long-term potentiation and increases in thiobarbituric acid reactive substances (TBARS) content, which indicate oxidative stress on hippocampal neurons responsible for cognitive impairments (Alhowail, Bloemer et al. 2019). Higher loads of oxidative species also target differentiating oligodendrocytes progenitor cells and myelin producing oligodendrocytes which eventually influence sensorimotor function (Brown, Stemmer et al. 1998, Maas, Valles et al. 2017). Because Nrf2 is an endogenous master regulator of antioxidant genes, it is an attractive therapeutic target to restore redox homeostasis. Fumarates effectively activate Nrf2 in clinical and preclinical studies (Linker, Lee et al. 2011, Hoogendoorn, Avery et al. 2021). Dimethyl fumarate (DMF), DRF, and MMF are all FDA approved to treat multiple sclerosis and psoriasis (Schimrigk, Brune et al. 2006, Linker, Lee et al. 2011, Carlstrom, Ewing et al. 2019, Cuadrado, Rojo et al. 2019). Nrf2 activators are also neuroprotective in neurodegenerative disease models. Pre-clinical studies in rodent model of Alzheimer’s disease show that pharmacological activation of Nrf2 by DMF prevent the deficit in spatial and working memory, and the associated neurotoxicity (Majkutewicz, Kurowska et al. 2016). In Oxidative stress in the substantia nigra pars (SNpc)-striatum neuroaxis is thought to be one of the cause of dopaminergic neurodegeneration in Parkinson’s disease (PD). Such patients exhibit increased levels of oxidized lipids and decreased levels of reduced glutathione (GSH) in the SNpc (Bosco, Fowler et al. 2006, Zeevalk, Razmpour et al. 2008). DMF has been shown to protect mouse nigrostriatal neurons and SHSY-5Y cells by augmenting Nrf2-mediated antioxidant pathways after 1-methyl-4-phenyl-1,2,3,6-tetrahydropyridine (MPTP) exposure (Ahuja, Ammal Kaidery et al. 2016, Campolo, Casili et al. 2017). Further, Nrf2 deficient mice show an increased level of proinflammatory markers such as cyclooxygenase-2 (COX-2), inducible nitric oxide synthase (iNOS), interleukin-6 (IL-6), and tumor necrosis factor-α (TNF-α) in MPTP-induced PD model which is indicative of microgliosis (Rojo, Innamorato et al. 2010).

Nrf2 expression is not limited to the brain, with systemic activators potentially interacting with Nrf2 globally causing unwanted effects. A clinical study of the systemic Nrf2 activator bardoxolone methyl administration to the patients with type 2 diabetes and stage-4 chronic kidney disease found increased risk of death from cardiovascular abnormalities (de Zeeuw, Akizawa et al. 2013). Nrf2 activation by oltipraz in pre-clinical rodent model induced systemic hypertension and kidney injury (Zhao, Ghosh et al. 2018). Activation of Nrf2 may have a role in carcinogenesis (Dinkova-Kostova and Copple 2023). Notably, activation of Nrf2 during carcinogenesis generates transcription enhancers at gene loci different than transiently activated Nrf2 (Okazaki, Anzawa et al. 2020). The electrophilic agent MMF succinates thiol groups of Kelch like ECH-associated protein 1 (Keap1)-Nrf2 complex in cytosol. This then releases Nrf2 from the complex to enter the nucleus. DRF systemically releases MMF interact with protein thiols other than those of Keap1. These off target interactions and associated adverse effects can be avoided with prodrug **1c** which releases MMF only in tissues with pathological concentration of oxidative species (Avery, Li et al. 2021). In this study, we administered DRF or **1c** to reverse doxorubicin-induced toxicities in mice, which were similarly effective.

Collectively, our findings demonstrate that both global and local Nrf2 activators reverse doxorubicin-induced cognitive impairments and sensorimotor deficits in WT mice of both sexes. The behavioral impairments were associated with loss of synaptic protein, morphological changes in microglia in the CA1 region of the hippocampus, and myelin damage in the corpus callosum. These effects are likely driven by oxidative stress, since they were all rescued by DRF and **1c**. Our study indicates that diroximel fumarate and **1c** should be further explored to treat the growing number of cancer survivors who suffer from long-lasting neurotoxic side effects of doxorubicin.

## Acknowledgments

This work was supported by National Institutes of Health grants RF1NS113840 (P. Grace), UG3NS127251 (A. Abell, P. Grace), and in part by P30CA016672 (animal housing and care in the MD Anderson Research Animal Support Facility, EM in the High-Resolution Electron Microscopy Facility); and Australian Research Council grants CE140100003 and DP180101581 (A. Abell). We thank Mr. Kenneth Dunner, Jr., High Resolution Electron Microscopy Facility (EM Core) for his excellent technical assistance for sample preparation and TEM imaging.

---

**Fig. S1:**
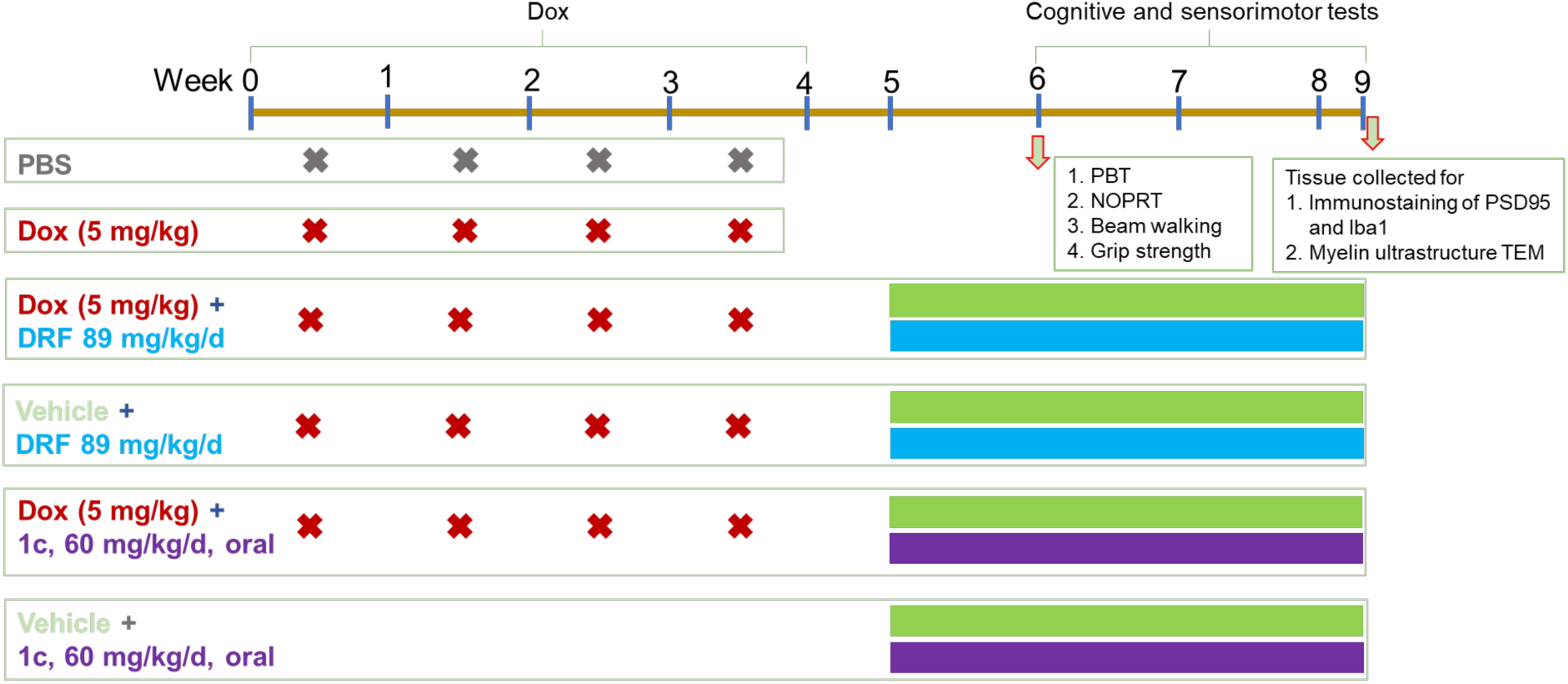
Dosing schedule and experimental design. Male and female mice were treated with PBS or doxorubicin (5 mg/kg), intraperitoneal, once a week for 4 consecutive weeks. DRF (89 mg/kg, oral) or **1c** (60 mg/kg, oral) was administered daily, starting one week after doxorubicin treatment and continuing through out the cognitive and sensorimotor tests. Cognitive tests (PBT and NOPRT) were started 2 weeks after the last dose of doxorubicin and followed by sensorimotor tests. Tissue was collected for TEM and immunostaining after completion of the tests.

